# Deep learning segmentation of the nucleus basalis of Meynert on 3T MRI

**DOI:** 10.1101/2022.07.02.498556

**Authors:** Derek J. Doss, Graham W. Johnson, Saramati Narasimhan, Jasmine W. Jiang, Hernán F. J. González, Danika L. Paulo, Alfredo Lucas, Kathryn A. Davis, Catie Chang, Victoria L. Morgan, Christos Constantinidis, Benoit M. Dawant, Dario J. Englot

## Abstract

The nucleus basalis of Meynert (NBM) is a key subcortical structure that is important in arousal, cognition, brain network modulation, and has been explored as a deep brain stimulation target. It has also been implicated in several disease states, including Alzheimer’s disease, Parkinson’s disease, and temporal lobe epilepsy (TLE). Given the small size of NBM and variability between patients, NBM is difficult to study; thus, accurate, patient-specific segmentation is needed. We investigated whether a deep learning network could produce accurate, patient-specific segmentations of NBM on commonly utilized 3T MRI. It is difficult to accurately segment NBM on 3T MRI, with 7T being preferred. Paired 3T and 7T MRI datasets of 21 healthy subjects were obtained, with 6 completely withheld for testing. NBM was expertly segmented on 7T MRI, providing accurate labels for the paired 3T MRI. An external dataset of 14 patients with TLE was used to test the model on brains with neurological disorders. A 3D-Unet convolutional neural network was constructed, and a 5-fold cross-validation was performed. The model was evaluated on healthy subjects using the held-out test dataset and the external dataset of TLE patients. The model demonstrated significantly improved dice coefficient over the standard probabilistic atlas for both healthy subjects (0.68MEAN±0.08SD vs. 0.47±0.06, p=0.0089, t-test) and TLE patients (0.63±0.08 vs. 0.38±0.19, p=0.0001). Additionally, the centroid distance was significantly decreased when using the model in patients with TLE (1.22±0.33mm, 3.25±2.57mm, p=0.0110). We developed the first model, to our knowledge, for automatic and accurate patient-specific segmentation of the NBM.

## Introduction

The nucleus basalis of Meynert (NBM) is a basal forebrain nucleus and is one of the major sources of cholinergic signal in the brain [2-4]. It has been implicated as abnormal in several disorders with cognitive decline such as Parkinson’s disease dementia, Alzheimer’s disease, and temporal lobe epilepsy (TLE) [5-7]. In Parkinson’s disease dementia, changes in the NBM predict the development of cognitive impairment [7]. Neuronal loss and atrophy of the NBM have both been observed in Alzheimer’s disease [8]. In TLE, the NBM has been shown to have decreased functional connectivity [6]. Additionally, there has recently been increased interest in studying the functional and structural network abnormalities involving NBM connections in several disease states [6, 9, 10]. However, there are several challenges to studying NBM anatomy.

While the NBM is thought to be important in several disease states, it is difficult to study due to pathological changes in the nucleus and pathological changes in disease states, such as atrophy [11]. It is possible to accurately segment the NBM, thus accounting for the patient-specific changes, but it is difficult to visualize the nucleus on commonly utilized 3T T1-weighted magnetic resonance imaging (MRI). More accurate manual segmentations can be performed on high-resolution 7T MRI (Fig. 1). To overcome this limitation, previous groups have developed a probabilistic atlas of the NBM [12]. This atlas was derived from histological slices of 10 healthy subject’s post-mortem. The segmented histological slices were then aligned to MNI152 space to provide a probabilistic atlas of the NBM. The probabilistic atlas has gained popularity and is the most commonly used atlas of the NBM. It has been implemented into the SPM anatomy toolbox for ease of use [13]. While the probabilistic atlas provides an accurate segmentation for most cases, it does not capture patient specific anatomical variability (Fig. 2). Similar problems in creating patient-specific segmentation of the NBM have been observed in other small brain nuclei [14-17]. Furthermore, recent studies have demonstrated the inconsistencies in using probabilistic atlas based segmentation of the NBM [18]. This challenge in particular has been recognized as one of the key limitations for studying the NBM in-vivo [18].

**Fig. 1:**
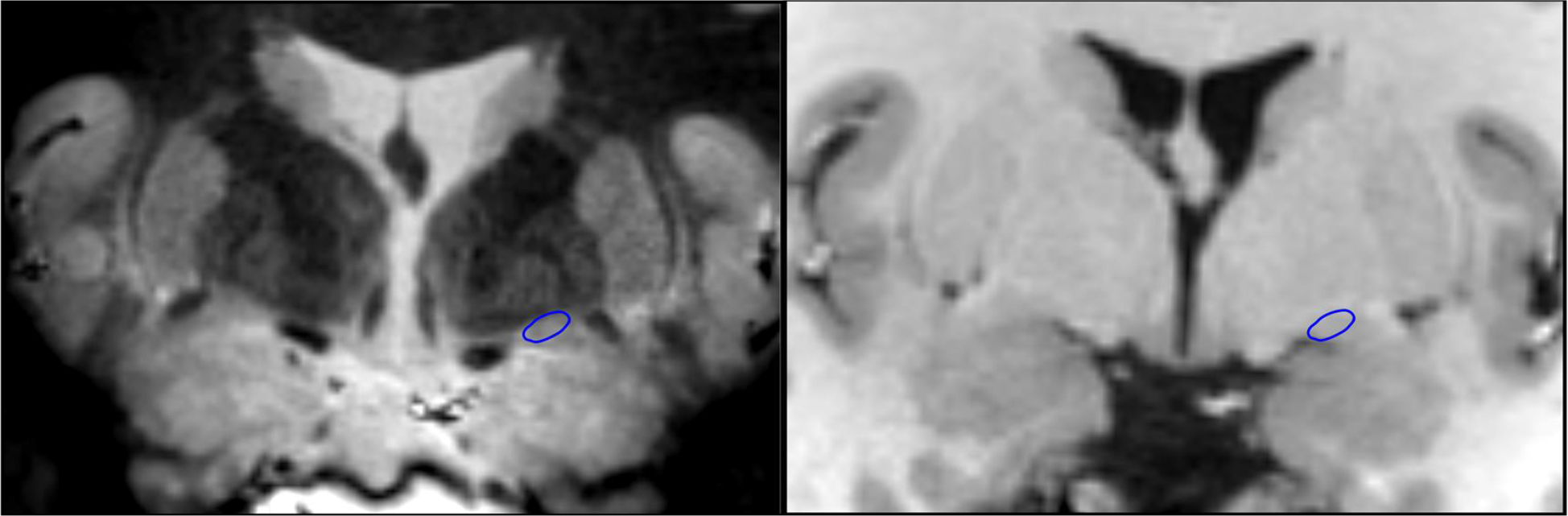
Similar coronal slice on the 7T MP-RAGE (left) and 3T T1-weighted (right) MRI scans of the same subject. The NBM is outlined in blue unilaterally and is more clearly visualized at 7T than 3T

**Fig. 2:**
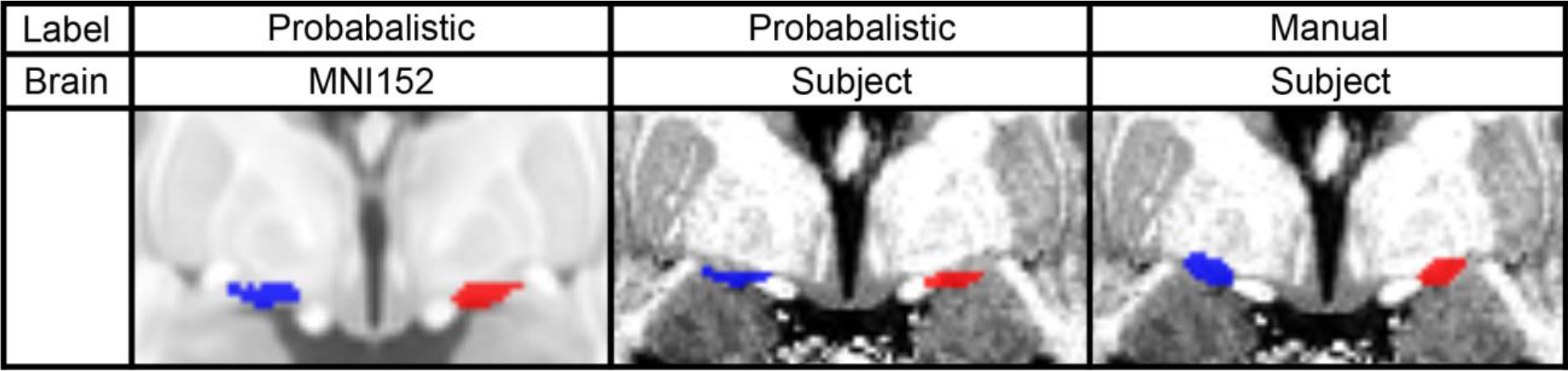
Comparison of the probabilistic and manual labels on a similar coronal slice of the MNI152 brain and a subject’s 7T brain. The left NBM is shown in blue and the right NBM is shown in red.

Thus, to overcome the above challenges, we propose a novel method for patient-specific NBM segmentation at 3T using subjects with both 3T and 7T MRI scans. The 3T scan can be used to train the deep learning network and NBM labels expertly segmented from 7T scans. Through this process, we can create a deep nuclei segmentation network (DnSeg) to produce an accurate, patient-specific segmentation of the NBM. The code and trained network are freely available at: https://github.com/DerekDoss/DnSeg.

## Methods

### Dataset

Both 7T and 3T MRI scans of the same healthy subjects were obtained. Data from a total of 21 healthy subjects were available. The entire dataset included data from two institutions, 11 subjects from Vanderbilt University and 10 subjects from University of Amsterdam [19-21]. The voxel sizes of the 3T data were 1.00mm x 1.00mm x 1.00mm. The voxel sizes of the 7T data were 0.70mm x 0.70mm x 0.70mm from Vanderbilt and 0.80mm x 0.80mm x 0.80mm from Amsterdam. A total of 6 healthy subjects from this dataset were completely withheld as a test dataset.

An additional external dataset of 14 paired 3T and 7T MRI of patients with TLE was obtained. Half of the patients had lesional changes on MRI and the other half were nonlesional. The voxel sizes of the 3T data were 0.80mm x 0.80mm x 0.80mm. The voxel sizes of the 7T data were 1.00mm x 0.95mm x 0.95mm. This dataset was withheld from the training and validation process and used only for testing of the deep learning network on brains with neurological disorders. The demographics of both datasets can be seen in Table I.

**TABLE I.**
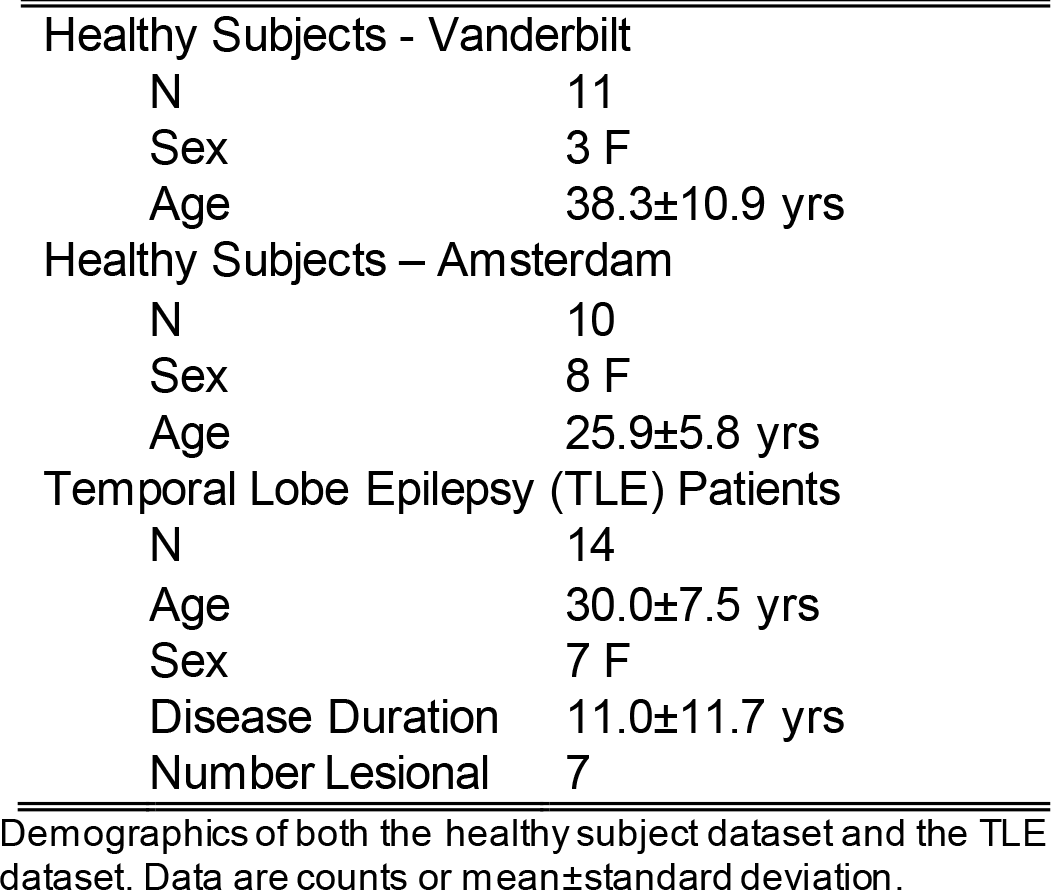
Dataset Demographics

|  |  |
| --- | --- |
| Healthy Subjects - Vanderbilt |  |
| N | 11 |
| Sex | 3 F |
| Age | 38.3±10.9 yrs |
| Healthy Subjects – Amsterdam |  |
| N | 10 |
| Sex | 8 F |
| Age | 25.9±5.8 yrs |
| Temporal Lobe Epilepsy (TLE) Patients |  |
| N | 14 |
| Age | 30.0±7.5 yrs |
| Sex | 7 F |
| Disease Duration | 11.0±11.7 yrs |
| Number Lesional | 7 |

Demographics of both the healthy subject dataset and the TLE dataset. Data are counts or mean±standard deviation.

### Manual Segmentation

Although the anatomical borders of the NBM cannot be easily discerned on 3T MRI, 7T MRI enables enhanced contrast in the NBM. According to the landmarks described in the literature, the anatomical borders of the NBM were identified [22, 23]. The anterior border was the anterior commissure, posterior border was the mamillary bodies, inferior border was the base of the basal forebrain, lateral border was the lateral portion of the anterior commissure, and medial border was the optic nerve. The left and right NBM were segmented for each subject and patient using our in house CRAnial Vault Explorer (CRAVE) software [1]. The segmentations of the NBM were completed by one author and were verified by two neurological surgeons.

### Data Preprocessing

The 7T and 3T paired images were converted to RAS orientation using nibabel [24]. The whole brain was extracted from the MRI scans using robust brain extraction (ROBEX) [25]. The 3T scan from each subject was rigidly registered to their 7T scan using SPM8 (https://www.fil.ion.ucl.ac.uk/spm/software/spm8/) (Fig. 3). The registration was manually verified to ensure that the NBM segmentation was accurate on the 3T scan. Afterwards the NBM segmentation was resliced to 1.0mm x 1.0mm x 1.0mm.

**Fig. 3:**
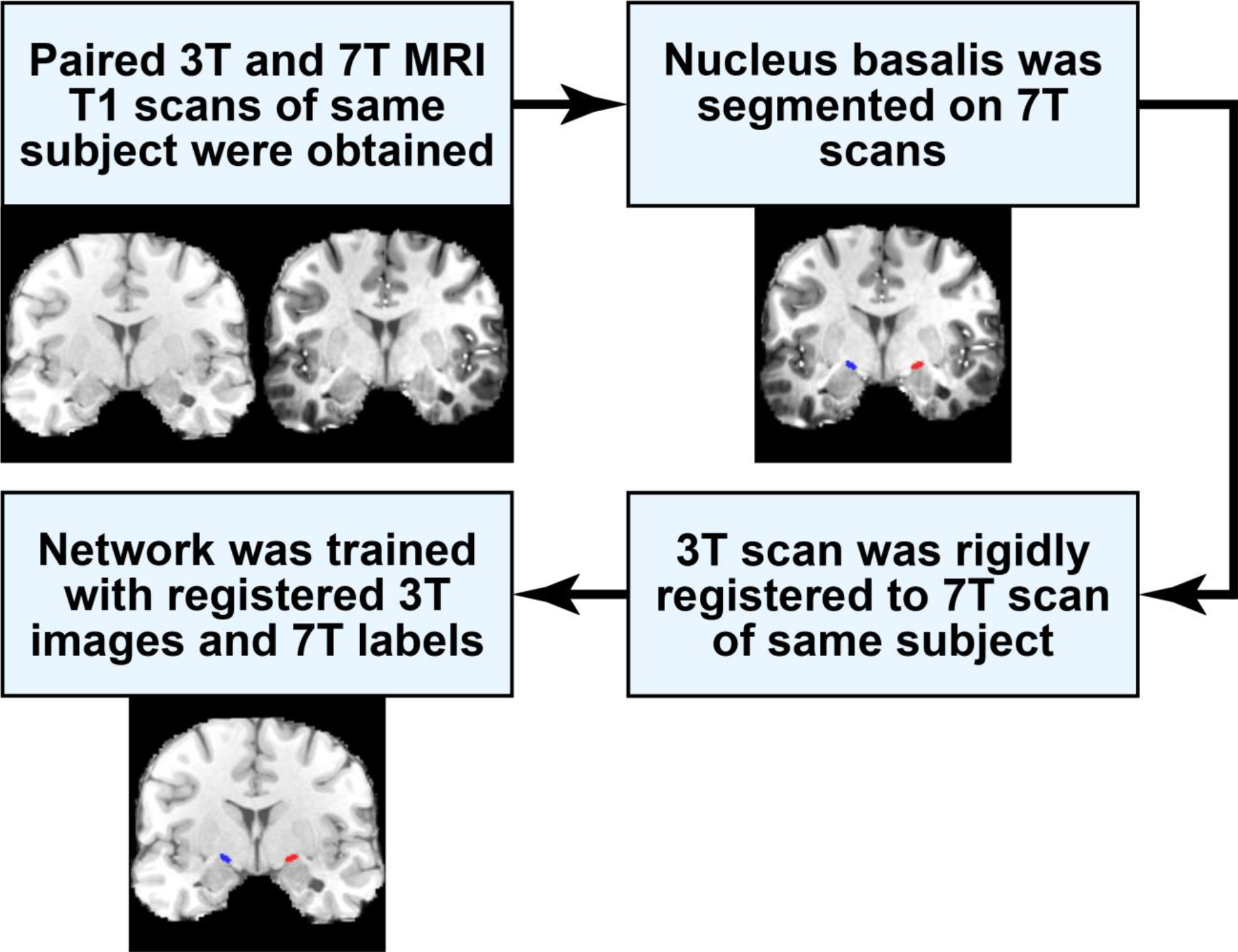
Flowchart describing how the 7T NBM segmentation is used for training the network to segment the NBM on 3T MRI.

### Data Augmentation

To enhance model generalization, we augmented the healthy subject dataset from 21 to 210 samples. Furthermore, aggressive augmentation allows for robust learning of anatomical variability. Augmentations performed include random rigid affine transformations, random elastic transformations, and random bias field additions (Fig. 4). To account for differing rotations that a subject’s head may experience in the MRI scanner, random affine transformations (rotations only) ranging from −10° to 10° in the x, y, and z axes were applied. To account for artifacts that can be present in MRI scanners due to a non-uniform magnetic field, a bias field of random intensity and direction was introduced into the images. A random elastic deformation field was also applied to the datasets, to ensure that the network was trained with substantial anatomic variability. The elastic deformation field had a max displacement of 30mm and 5 control points. All augmentations were performed using the TorchIO python library [26].

**Fig. 4:**
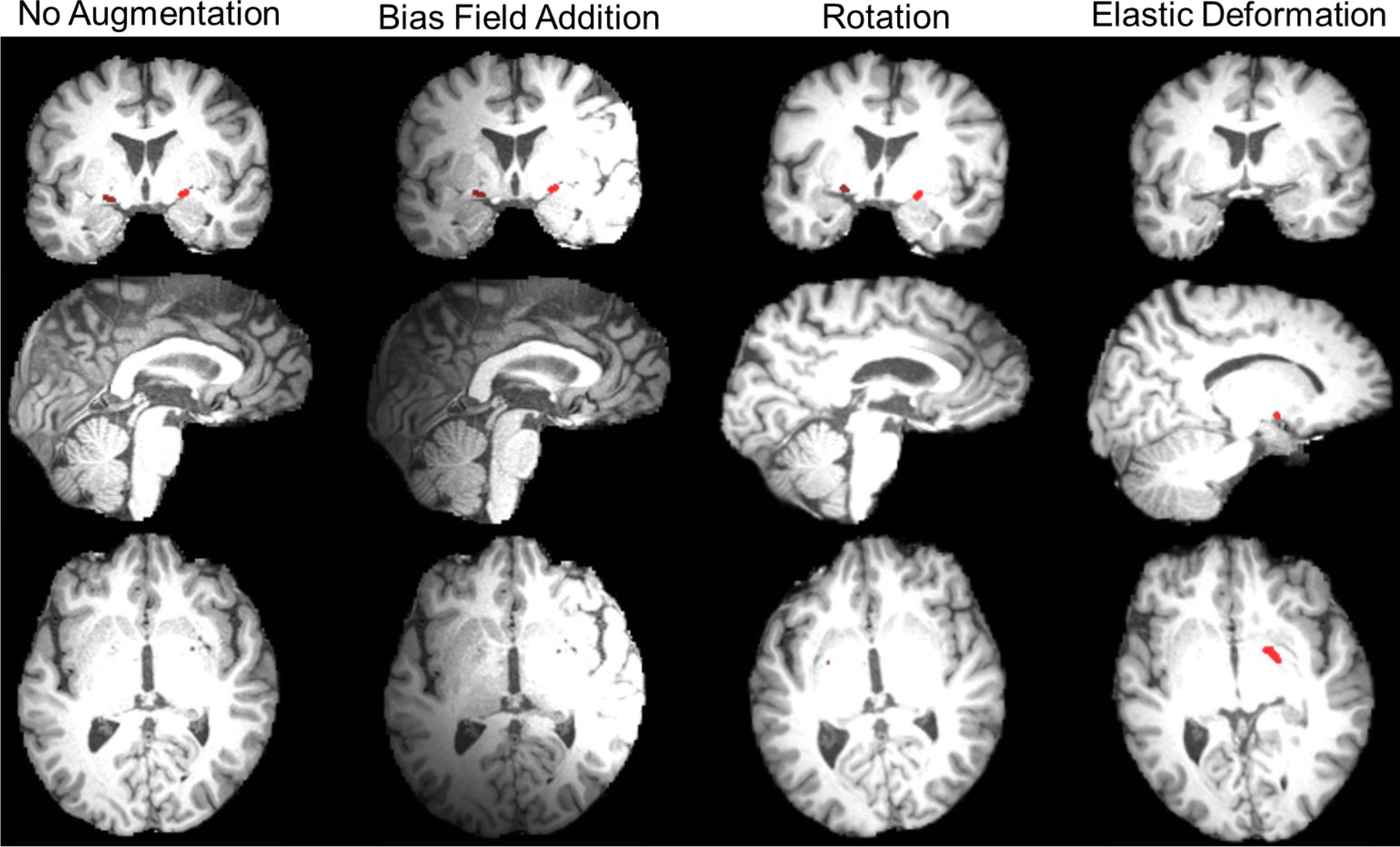
The same coordinates in a subject’s 3T MRI with various data augmentations performed. The left NBM is dark red and the right NBM is light red.

### Volume Reduction

Considering the large amount of video RAM required to train a deep learning network using the whole brain, the size of the images was reduced for ease of training. Given that the mamillary bodies are the posterior anatomical borders of the NBM, we used the patient specific segmentation package implemented into FreeSurfer ScLimbic to segment the mamillary bodies [16]. The mamillary bodies were used as the center of a 64mm x 64mm x 64mm region, which includes the entire NBM in all cases.

### Convolutional Neural Network

The U-net architecture for biomedical image segmentation was modified to accommodate 3D image segmentation [27]. A complete visualization of the architecture can be seen in Fig. 5. The network consisted of two convolutions with a kernel size of 5, padding of 1, and a stride of 2. The double convolution was followed by a max pool step. A total of 3 layers were implemented in the final model. A batch size of 1 was used. The network was trained using the soft dice loss function.

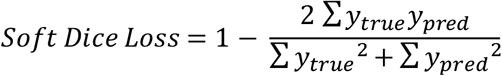

The data were divided into a train, test, and validation set. The datasets were split on the subject level. The train dataset consisted of 12 subjects (120 scans), the validation dataset consisted of 3 subjects (30 scans), and the test dataset consisted of 6 subjects (60 scans). The test dataset was completely held out from model selection and only used for the evaluation after the final model was selected.

A 5-fold cross validation was utilized on the train and validation datasets. The average dice performance from each fold on the validation dataset was utilized as the performance measure for model selection. Hyperparameter tuning during model selection was determined using a Bayesian hyperparameter search algorithm, hyperopt [28].

**Fig. 5:**
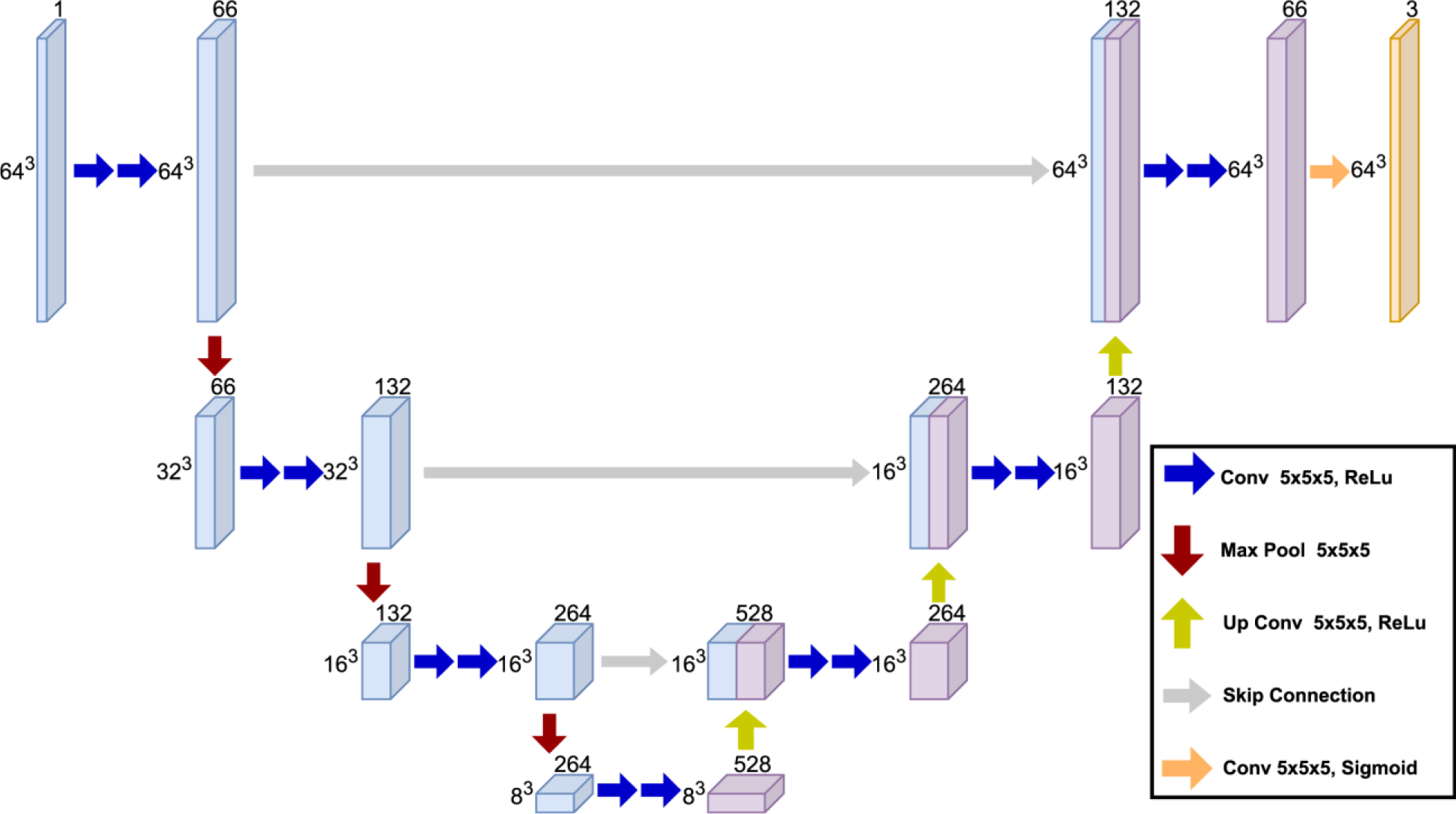
Visualization of the 3D-Unet created.

### Evaluation Metrics

Evaluation of DnSeg encompassed methods to assess volumetric overlap of the NBM and localization of the NBM. Conventional “hard” dice coefficient was used to determine the overlap between the predicted and manually segmented NBM [29].

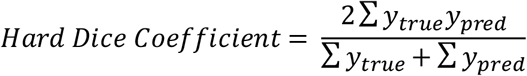

Mean surface distance was defined as the mean distance from the surface of the predicted and ground truth mesh [14, 15]. Centroid distance was defined as the Euclidean distance from the centroid of the predicted NBM and the ground truth segmentation [14, 15]. Centroid distance is relevant for the NBM as it more accurately represents targeting error that can occur during stereotactic neurosurgery.

The performance of DnSeg was compared against that of the probabilistic atlas. The probabilistic atlas was thresholded such that all voxels with a >50% probability of being NBM were assigned as NBM. The probabilistic atlas was registered from MNI152 to patient specific space using SPM8. The same metrics were then computed on the probabilistic atlas segmentation of the NBM.

## Results

All results were computed in patient 3T MRI space with 1mm isotropic slices. The results were all computed on the test dataset of 6 healthy subjects (60 augmented scans) which were held out from the training and validation of the DnSeg. Results were also computed on a held-out external dataset of 14 subjects with diagnosed TLE that were not used in development of the DnSeg.

### Dice Coefficient

Quantitative evaluation of the overlap of the predicted and ground truth segmented NBM was completed with the dice coefficient [29]. Dice coefficient is known to underestimate the performance of small, complex structures and the performance of DnSeg was similar to other small structure segmentation methods [14-17, 30, 31]. The Dice coefficient of the NBM computed from DnSeg (0.68MEAN±0.08SD) significantly outperformed that of the probabilistic atlas (0.47±0.06) in the completely withheld dataset of 6 healthy subjects (paired t-test p=0.0089).

The Dice coefficient was also computed for the dataset of patients with TLE, as can be seen in Fig. 6. The performance of the probabilistic atlas and that of DnSeg was compared. Of note, there was no significant difference between the Dice coefficient of DnSeg on the healthy subject and TLE dataset. However, there was a significant difference between DnSeg (0.63±0.08) and probabilistic atlas dice coefficient (0.38±0.19) for the TLE dataset (paired t-test p=0.0001). The probabilistic atlas demonstrated substantial variability in Dice coefficient for the TLE dataset, most likely due to the anatomical changes that have been noted in the NBM for TLE.

**Fig. 6:**
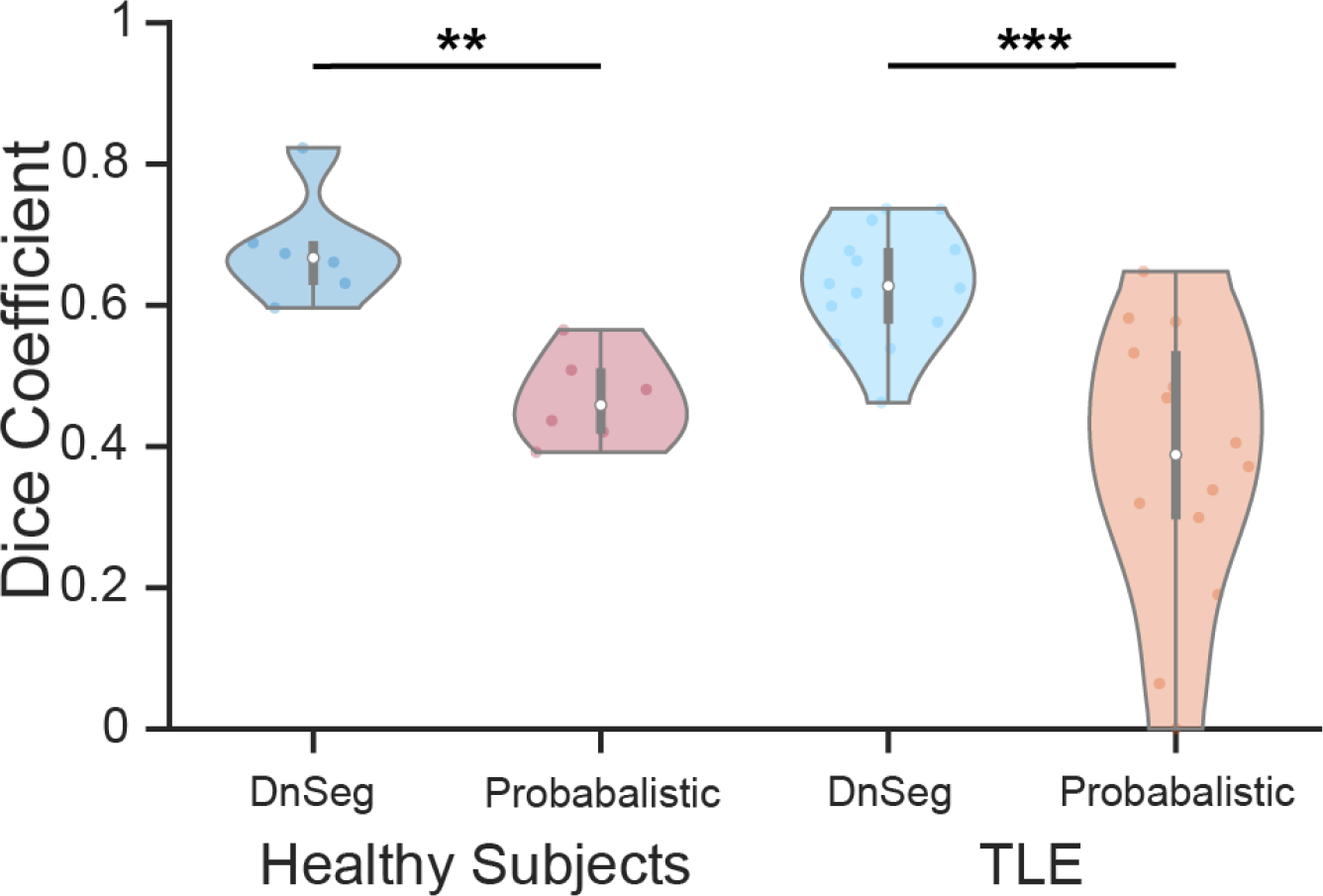
Dice coefficient of DnSeg vs the probabilistic atlas for both healthy subjects and patients with TLE. The healthy subjects included the held-out test dataset (N=6). The patients with TLE were also held-out until final analysis (N=14). Paired t-test: *p<0.05, **p<0.01, ***p<0.001

### Mean Surface Distance

Mean surface distance was used to estimate how closely the predicted segmentation edges matched the ground truth. The predicted surface and the ground truth surface of the NBM segmentation were calculated for the test dataset and the TLE dataset. The results can be seen in Fig. 7. The mean surface distance of the healthy subjects computed with DnSeg (0.67±0.18 mm) was significantly lower than the mean surface distance computed with the probabilistic atlas (1.14±0.18 mm) in the completely withheld dataset of 6 healthy subjects (paired t-test p=0.0090). Furthermore, the mean surface distance was significantly lower with DnSeg (0.72±0.12 mm) compared to the probabilistic atlas (2.00±2.11 mm) in the TLE dataset (paired t-test p=0.0381). The variability of the probabilistic atlas on the TLE dataset was far more exaggerated than that of the healthy subject dataset.

**Fig. 7:**
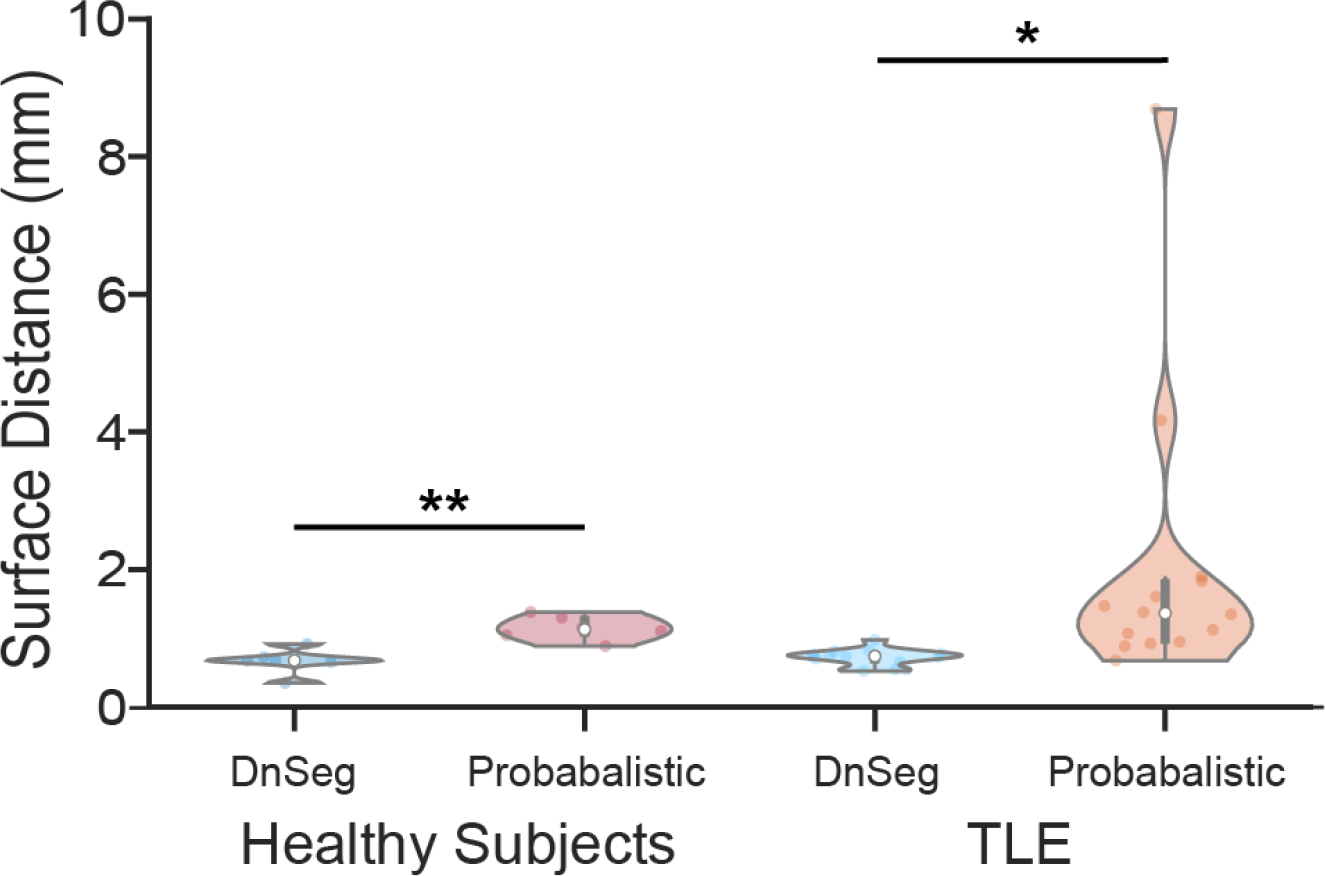
Average surface distance of the DnSeg vs the probabilistic atlas for both healthy subjects and patients with TLE. The healthy subjects includ ed the held-out test dataset (N=6). The patients with TLE were also held-out until final analysis (N=14). Paired t-test: *p<0.05, **p<0.01, ***p<0.001

### Centroid Distance

The centroid distance represents the error in localization of the NBM that can occur due to patient-specific anatomical variability. It is most relevant for possible DBS applications, as it could contribute to targeting error. The results of this analysis can be seen in Fig. 8. The centroid distance performance of DnSeg (1.41±0.73 mm) was not significantly different from the probabilistic atlas (2.23±0.85 mm) in the completely withheld dataset of 6 healthy subjects (paired t-test p=0.0510). However, the centroid distance of DnSeg (1.22±0.33 mm) was significantly lower than that of the probabilistic atlas (3.25±2.57 mm) in the TLE dataset (paired t-test p=0.0110).

**Fig. 8:**
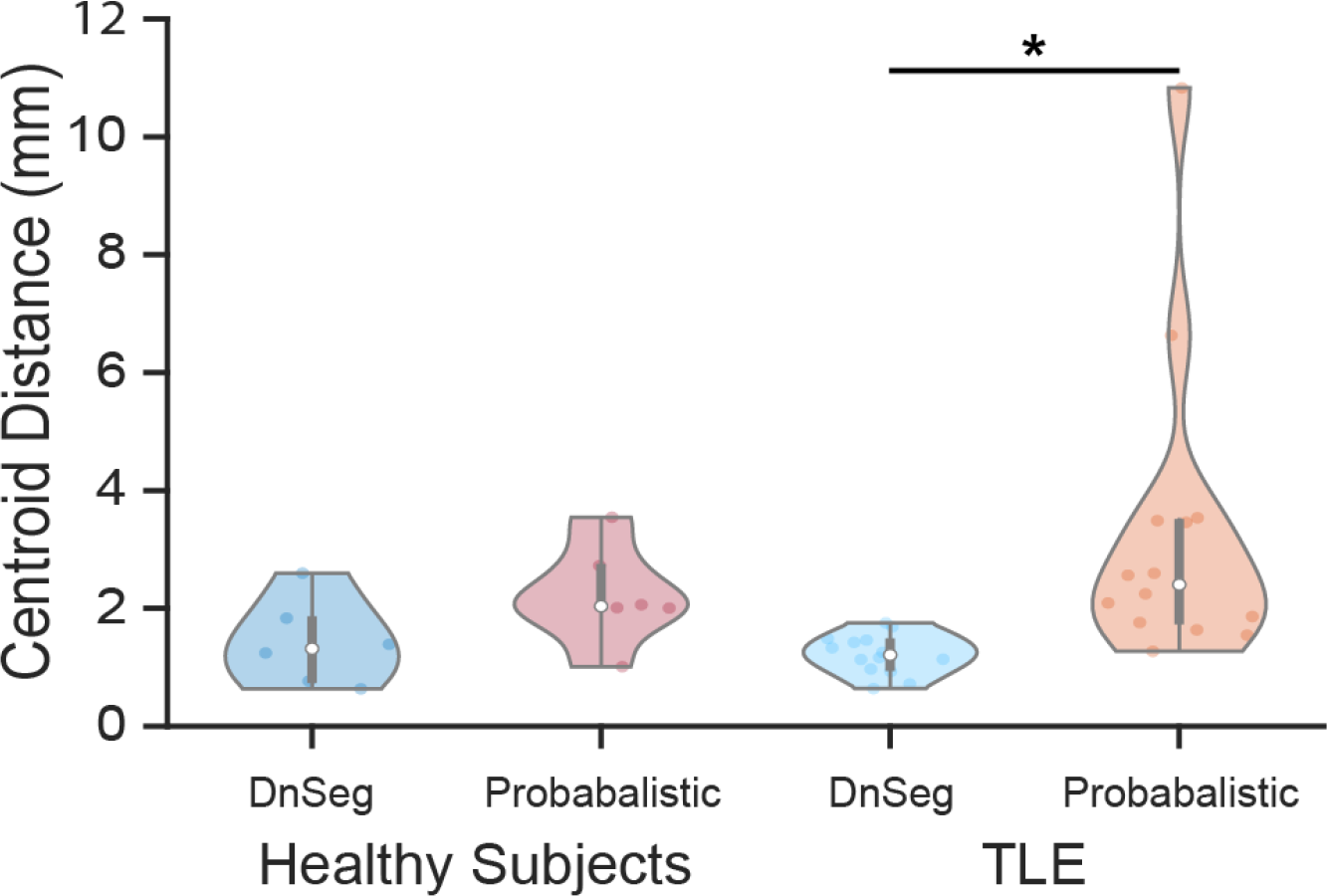
Centroid distance of DnSeg vs the probabilistic atlas for both healthy subjects and patients with TLE. The healthy subjects included the held-out test dataset (N=6). The patients with TLE were also held-out until final analysis (N=14). Paired t-test: *p<0.05, **p<0.01, ***p<0.001

### Qualitative Comparison

An evaluation of DnSeg’s NBM prediction compared to the ground truth manual segmentation was completed qualitatively by visual inspection. Fig. 9 shows this comparison. As can be seen, the average prediction error is around an extra voxel of over and under prediction on either side of the NBM. The poor performance example demonstrates that even the poor performing examples perform relatively well. Additionally, it is seen that the NBM shape differs slightly from patient to patient and that DnSeg successfully accounts for heterogenous patient anatomy.

**Fig. 9:**
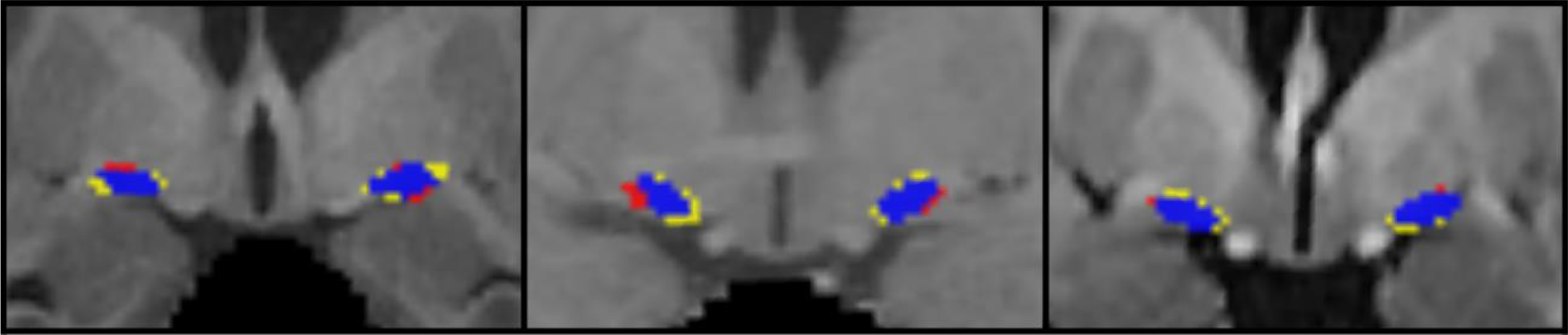
Example of favorable (right), average (middle), and poor (left) performance. Blue is the correct prediction of NBM, red is the over predicted NBM (false positive), and yellow is the ground truth label that was not predicted (false negative). The Dice coefficient of the favorable example (right) was 0.82, the Dice coefficient of the average example (middle) was 0.63, and the Dice coefficient of the poor example (left) was 0.55.

## Discussion

This work presents, to our knowledge, the first deep learning framework for segmentation of the NBM. The NBM is a challenging brain structure to segment as it cannot be accurately visualized on 3T MRI, and it is a small structure. Since it can be visualized accurately on 7T MRI, a novel approach was taken in which the NBM was segmented on 7T MRI and a deep neural network was trained to perform the segmentation on registered 3T MRI of the same subject, allowing for accurate NBM segmentation on commonly utilized 3T MRI.

### DnSeg can distinguish subject specific anatomical differences

DnSeg was first evaluated with dice coefficient; however, small structures have an artificially deflated performance with the dice coefficient. Although a dice coefficient of 0.68 is not ideal, our results are comparable to those of other small structure segmentation frameworks, such as those that segment the interposed nuclei or ventral tegmental area, which have resulted in dice coefficients of 0.69 and 0.56 respectively [15, 31].

The NBM has anatomical variability between healthy subjects and has been shown to change in size in patients with Parkinson’s disease, Alzheimer’s disease, and TLE [32-40]. It is difficult to capture this anatomical variability with manual segmentation of commonly used 3T T1-weighted MRI as the NBM cannot be accurately visualized on 3T MRI. The probabilistic atlas sought to solve the segmentation challenge; however, it seems to not capture patient-specific anatomical differences in disease states. Across multiple performance metrics (Dice coefficient, centroid distance, and average surface distance), the probabilistic atlas had wide variance in the TLE dataset and much smaller variance in the healthy subject test dataset. This likely indicates that while the probabilistic atlas can approximately capture the NBM in most healthy subjects, it is unable to capture the anatomical variability in patients. Given that the disease states of interest for studying the NBM are associated with changes in the neuronal count, shape, and size of the NBM, it is important for an atlas to capture the patient-specific anatomical variability. Based on the presented data, DnSeg accomplishes this goal.

The ability of DnSeg to distinguish subject-specific anatomy is shown in healthy subjects as well, as seen in Fig. 1. Since the anatomical variability in healthy subjects is less substantial than that of patients with TLE, the difference in performance of DnSeg and probabilistic atlas is less substantial yet still significant. These results demonstrate the ability of DnSeg to segment structures whose contrast differences are not readily apparent.

### DBS Targeting

DBS targeting for the NBM has been explored in several studies with mixed results [41-45]. There are several factors contributing to surgical success or failure, but one factor critical to success of DBS is targeting accuracy. As can be seen in this investigation, the standard probabilistic atlas in patients with TLE has a rather large center of mass distance of 3.2mm. It is possible that the mixed results in other studies could be due to difficulties in targeting the NBM. This can be most easily observed in Fig. 10, which shows an electrode planned with DnSeg seems to target the NBM accurately while an electrode planned with the probabilistic atlas targets the border of the true NBM. This work provides a method to segment the NBM that performs well in at least one disease state in which the NBM may be functionally and anatomically altered.

**Fig. 10:**
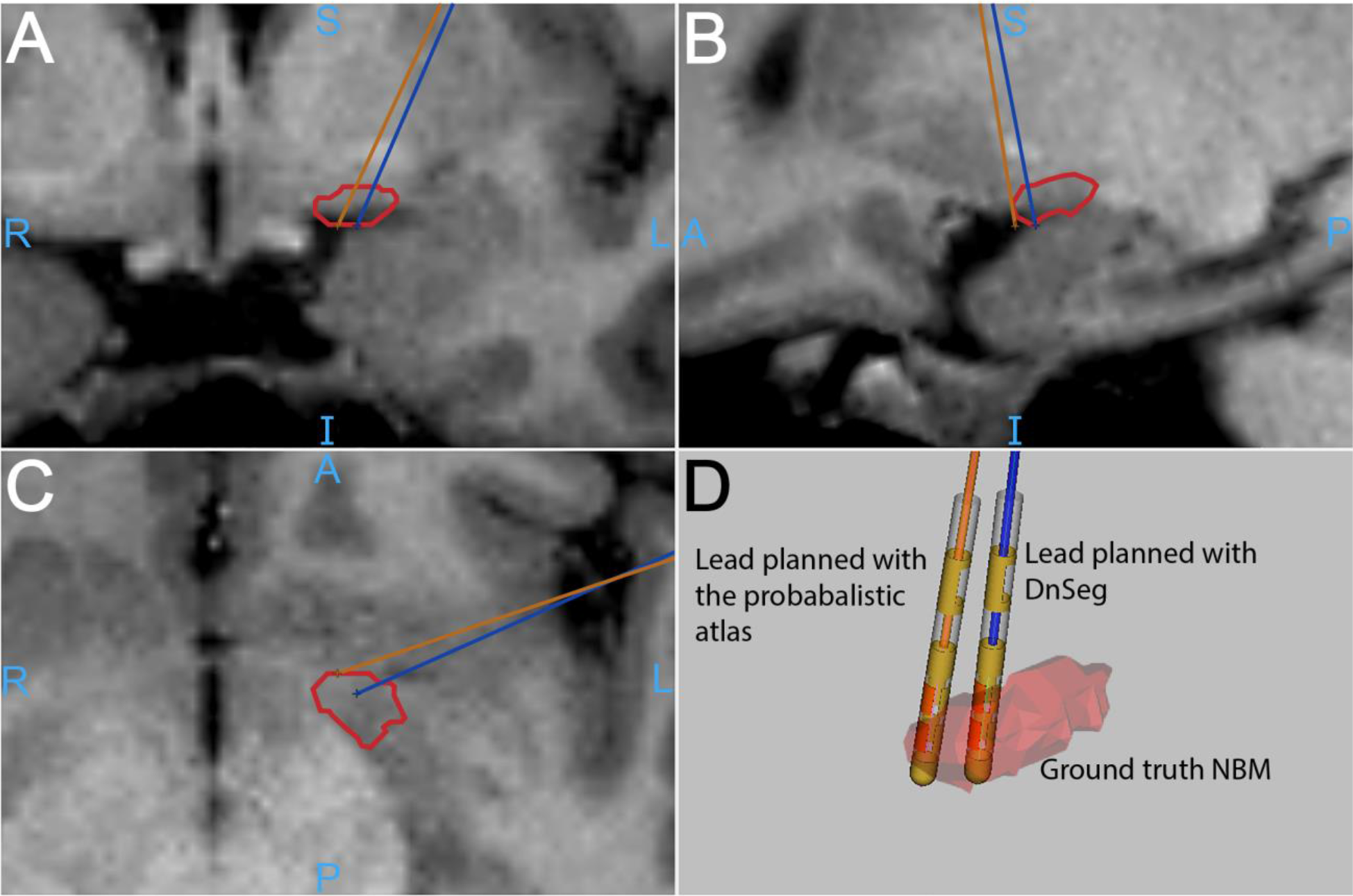
Increased accuracy of NBM electrode targeting with DnSeg. The orange electrode was placed in the center of the NBM segmentation generated by the probabilistic atlas. The blue electrode was placed in the center of the segmentation generated by DnSeg. Panel D shows both electrode positions compared to the ground truth segmentation mesh. The ground truth NBM (manually segmented on 7T MRI) is shown in red. The outline of the ground truth NBM as well as the lead trajectories can be seen in the panels A, B, and C. The example DBS planning for this figure was generated using CRAVE [1].

In addition to this practical application, DnSeg also has a much faster runtime than standard registration methods required for applying a MNI152 atlas to patient specific space. Running with a modern CPU, it takes approximately 1 minute per scan if skull stripping is not required and approximately 2 minutes if skull stripping is required. The pipeline does not require the patient data to be registered to any standard space, thus is much faster than other methods that require a registration step.

### Limitations

Despite these benefits of the deep learning pipeline, there are some limitations. Most notably, the expert segmentations could not be verified with a post-mortem histological analysis. While the segmentations were verified by two neurological surgeons, a histological analysis would be ideal. Furthermore, the current analysis only validates DnSeg in one disease state. The NBM is both of interest and has pathological changes in several disease states. Thus, it would be preferred to validate DnSeg in other disease states. However, for each patient, an expert segmentation is needed to evaluate the performance of DnSeg. As 7T MRI is preferred for accurate manual segmentation and 3T MRI is used by DnSeg to generate a predicted NBM segmentation, paired 7T-3T MRI scans of the same patients are needed. Therefore, the current analysis was limited by the rarity of paired 7T-3T MRI. For the described reason, DnSeg is also limited by the number of subjects used to train it. This has the potential to limit the generalizability of DnSeg. Aggressive augmentation was utilized to decrease the impact of this limitation and increase generalizability.

## Conclusions

In this work, we have presented an accurate, patient-specific method of segmenting the NBM. The NBM has been implicated in Alzheimer’s disease, Parkinson’s disease dementia, and TLE [6, 9, 10]. It has been shown to change in neuronal density and gray matter volume across age and disease states [32-40, 46]. The use of expertly segmented NBM and aggressive data augmentation, we have trained a deep learning network to capture anatomical differences when segmenting the NBM. We have presented evidence that DnSeg is a powerful and accurate NBM segmentation model.

Methodologically, this work represents an innovative approach to the segmentation of regions with little contrast enhancement. The NBM represents a small region of the brain with little contrast on commonly utilized 3T MRI and high importance to several disease states. Using of paired 3T and 7T MRI of the same subjects is a useful approach to providing accurate labels for training a deep learning network to segment structures that cannot be accurately visualized on 3T imaging. However, this approach is intrinsically limited by the rarity of paired 3T-7T datasets.

DnSeg, therefore, opens possibilities of further study of the NBM. The NBM has long been considered a region of key interest, but study of the NBM in-vivo has been difficult in part because of the lack of an adaptive, patient-specific atlas. The presented model is available (https://github.com/DerekDoss/DnSeg) and may greatly assist in novel studies of the NBM.

## Acknowledgment

The authors would like to thank Laurie Cutting, Ph.D. for providing paired 7T-3T MRI scans of healthy subjects. The authors would also like to thank William (Bill) Rodriguez, M.Ed. for his assistance with CRAVE and dataset curation.

## Notes

### Competing Interest Statement

The authors have declared no competing interest.

### Summary of Updates

The license was updated.

